# Transient receptor vanilloid 4 (TRPV4) channels are essential for alveolar epithelial cell function

**DOI:** 10.1101/775668

**Authors:** Jonas Weber, Yu-Kai Chao, Martina Kannler, Gabriela Krasteva-Christ, Suhasini Rajan, Ali Önder Yildirim, Monika Brosien, Johann Schredelseker, Norbert Weissmann, Christian Grimm, Thomas Gudermann, Alexander Dietrich

## Abstract

Ischemia-reperfusion(IR)-induced edema formation can be mimicked ex-vivo in isolated perfused mouse lungs (IPL). Here we show enhanced edema formation in transient receptor potential vanilloid 4 (TRPV4)-deficient (TRPV4-/-) IPL compared to wild-type (WT) controls in response to IR, indicating a protective role of TRPV4 to maintain the alveolar epithelial barrier. By immunohistochemistry, mRNA profiling or electrophysiological analysis we detected TRPV4 in bronchial epithelium, alveolar type I (ATI) and alveolar type II (ATII) cells. Genetic ablation of TRPV4 resulted in reduced expression of aquaporin-5 (AQP-5) channels in ATI as well as decreased production of pro surfactant protein C (pSP-C) in ATII cells. Migration of TRPV4-deficient ATI cells was reduced and cell barrier function was impaired. Moreover, adult TRPV4−/− lungs developed emphysema-like changes and altered lung parameters compared to WT lungs. Therefore, our data highlight novel essential functions of TRPV4 channels in alveolar epithelial cells and in the protection from edema formation.

**eLife digest:** Transient receptor potential vanilloid 4 (TRPV4) is a non-selective Ca^2+^ permeable cation channel expressed in lung endothelium where increased channel activity has been shown to compromise endothelial barrier function. In other tissues however, the channel maintains physiological cell barriers, e.g. in skin, the urogenital tract and the corneal epithelium. In tracheal epithelial cells TRPV4 channels regulate ciliar beat frequency and in alveolar epithelial cells TRPV4 activation by 4α-phorbol esters produced blebs and breaks in lung septa by unknown molecular mechanisms. To understand the channels role in lung function Weber et al. employed ex-vivo isolated perfused mouse lungs (IPL) to mimic ischemia-reperfusion-induced edema as one of the most common and significant causes of morbidity and mortality after lung transplantation in human patients. TRPV4-deficient (TRPV4−/−) IPL developed enhanced edema formation compared to wild-type (WT) controls in response to ischemia and reperfusion, indicating a protective role of TRPV4 to maintain the alveolar epithelial barrier. TRPV4 was detected in bronchial epithelium, alveolar type I (ATI) and alveolar type II (ATII) cells by immunohistochemistry or mRNA profiling. Genetic ablation of TRPV4 resulted in reduced expression and plasma membrane insertion of water conducting aquaporin-5 (AQP-5) channels in ATI cells compared to WT mice. Analysis of isolated primary TRPV4−/− ATII cells revealed a reduced expression of pro surfactant protein-C (pSP-C) a precursor of a protein important for decreasing surface tension and for alveolar fluid homeostasis. Moreover, the TRPV4 activator GSK1016790A induced increases in current densities only in WT but not in TRPV4−/− ATII cells. On a molecular level ablation of TRPV4 induced less Ca^2+^-mediated nuclear translocation of nuclear factor of activated T-cells (NFAT) to the nucleus, which may be responsible for reduced expression of the identified proteins. Although the ability of TRPV4−/− ATII to differentiate to ATI cells was unchanged, migration of TRPV4-deficient ATI cells was reduced and cell barrier function was impaired. Moreover, TRPV4−/− lungs of adult mice developed significantly larger mean chord lengths and altered lung function compared to WT lungs. The findings of Weber et al. highlights novel essential functions of TRPV4 channels in alveolar epithelial cells and in the protection from edema formation.

## Introduction

The alveolar epithelium serves multiple functions in the lung. On the one hand, the epithelial layer forms a natural barrier to the external environment protecting the body from invading microorganisms and toxicants, while, on the other hand, alveolar epithelial cells facilitate gas exchange. In the adult lung, the alveolar epithelium consists of two epithelial cell types, which are crucial to maintain lung homeostasis and tissue repair (*Mutze et al., 2015*). Alveolar epithelial type I (ATI) cells are elongated cells with a large cell surface and high barrier function in close proximity to endothelial cells of the alveolar capillaries facilitating gas exchange (*Mutze et al., 2015*). ATI cells are also highly water permeable, allowing for ion transport and maintenance of lung fluid balance (*Dobbs et al., 2010*). Although the latter cells cover the largest surface area of the lung (*Weibel, 2015*), alveolar epithelial type II (ATII) cells, which exhibit a cubic morphology, by far outnumber ATI cells (*Stone et al., 1992*). ATII cells are also involved in ion transport and liquid homeostasis *(Fehrenbach, 2001*) and are – most importantly - responsible for the production, storage, secretion and recycling of pulmonary surfactant. Surfactant lowers the surface tension at the tissue-air barrier to allow proper inflation and deflation of the lung during breathing (*Halliday, 2008*). Moreover, ATII cells also serve as progenitors for ATI cells and are capable of long-term self-renewal (*Desai et al., 2014*). Although alveolar epithelial cells express a wide variety of ion transport proteins and channels (*Hollenhorst et al., 2011*), their exact roles for specialized alveolar cell functions have still remained elusive.

Transient receptor potential vanilloid 4 (TRPV4) is the fourth cloned member of the vanilloid family of TRP channels (*Nilius and Szallasi, 2014*). Like most TRP channels TRPV4 harbors an invariant sequence, the TRP box (containing the amino acid sequence: EWKFAR), in its intracellular C-terminal tail as well as ankyrin repeats in the intracellular N-terminus. The protein is composed of six membrane-spanning helices (S1-6), and a presumed pore-forming loop between S5 and S6 *(Dietrich et al., 2017; Nilius and Szallasi, 2014*). Four of these monomers of the same type preferentially assemble in a functional homotetrameric complex (*Hellwig et al., 2005*), although in cell cilia of renal epithelial cells TRPV4/TRPP2 complexes were also identified (*Kottgen et al., 2008*). Homotetrameric TRPV4 was originally characterized as a sensor of extracellular osmolarity (*Liedtke et al., 2000; Strotmann et al., 2000*). The channel is functionally expressed in endothelial (*Hill-Eubanks et al., 2014*) and epithelial cells of the respiratory system (*Li et al., 2011; Lorenzo et al., 2008*). TRPV4 channels are thermosensitive in the range from 24 to 38 °C and may additionally serve as mechanosensors, because they are activated by membrane and shear stretch as well as by viscous loading (*Goldenberg et al., 2015a*). As TRPV4 is also involved in pulmonary hypertension (*Goldenberg et al., 2015b; Xia et al., 2013*) and bladder function (*Everaerts et al., 2010*), the channel is an interesting pharmacological target and numerous modulators have already been identified (reviewed in (*Dietrich, 2019*)). Moreover, TRPV4-deficient mice were protected from bleomycin-induced pulmonary fibrosis, due to the channel’s constitutive expression and function in lung fibroblasts (*Rahaman et al., 2014*). In lung endothelium, where its role was most extensively studied, direct or indirect activation of TRPV4 by mechanical stress (*Jian et al., 2008*), high peak inspiratory pressure (*Hamanaka et al., 2007; Michalick et al., 2017*) and high pulmonary venous pressure due to heart failure (*Thorneloe et al., 2012*) resulted in the disruption of the endothelial barrier and edema formation. In other tissues however, the channel maintains physiological cell barriers, e.g. in skin (*Akazawa et al., 2013*), the urogenital tract (*Janssen et al., 2016*) and the corneal epithelium (*Martinez-Rendon et al., 2017*). In tracheal epithelial cells TRPV4 channels regulate ciliar beat frequency (*Lorenzo et al., 2008*) and in alveolar epithelial cells TRPV4 activation by 4α-phorbol esters produced blebs and breaks in lung septa (*Alvarez et al., 2006*) by unknown molecular mechanisms. Moreover, stimulation of TRPV4 by bacterial lipopolysaccharides (LPS) mounted a protective response (*Alpizar et al., 2017*), while TRPV4 inhibition reduced lung edema and inflammation after chlorine exposure (*Balakrishna et al., 2014*). Therefore, TRPV4 channels may function as chemosensors of toxicants in the lung epithelium (reviewed in (*Steinritz et al., 2018*)).

In here, we set out to study pulmonary functions of TRPV4 channels capitalizing on a TRPV4-deficient mouse model. Triggered by ischemia-reperfusion, we observed enhanced lung edema formation, probably due to down-regulation of aquaporine-5 channels in alveolar type I (ATI) cells, reduced pro surfactant protein-C (pSP-C) production in ATII cells and/or emphysema-like changes in the overall lung architecture. Our data suggest an essential role of TRPV4 channels in the alveolar epithelium.

## Results

### Ablation of TRPV4 increases ischemia-reperfusion (IR)-induced edema formation in isolated perfused mouse lungs

To investigate the role of TRPV4 in IR-induced edema formation, we isolated lungs from wild-type (WT) and TRPV4-deficient (TRPV4−/−) mice. Initial characterization of these mice revealed impaired pressure sensation in dorsal root ganglia *(Suzuki et al., 2003*) and impaired osmotic sensation by exaggerated arginine vasopressin (AVP) secretion in the brain (*Mizuno et al., 2003*). Loss of TRPV4-protein was confirmed in lung lysates. While in WT controls a protein of the appropriate size of 100 kDa was detected by Western blotting with TRPV4 specific antibodies, TRPV4−/− lungs did not express any TRPV4 protein (*Figure 1A*). Murine embryonic fibroblasts (MEF) (*Kalwa et al., 2015*) like pulmonary fibroblasts express TRPV4 protein (*Rahaman et al., 2014*) and served as an additional positive control. After initial perfusion for 15 min followed by 90 min ischemia and 120 min reperfusion, TRPV4-deficient lungs show enhanced lung edema formation as evidenced by a considerable gain in lung weight as opposed to WT lungs (*Figure 1B*). Weight increased to a similar extent as already described by us previously (*Weissmann et al., 2012*). These results clearly contrast with observations on TRPC6-deficient lungs, which are protected from IR-induced edema due to reduced endothelial permeability (*Weissmann et al., 2012*). Therefore, we generated a TRPV4/TRPC6 double deficient (TRPV4/TRPC6−/−) mouse model, whose lungs lack the increase in IR-induced edema formation, but developed edema similar to WT mice (*Figure 1B*). Moreover, lung edema formation in TRPV4−/− lungs was clearly visible (*Figure 1C*) and consistently wet to dry weight ratio gain doubled in TRPV4−/−, but only slightly increased in TRPV4/TRPC6−/− lungs (*Figure 1D*).

**Figure 1.**
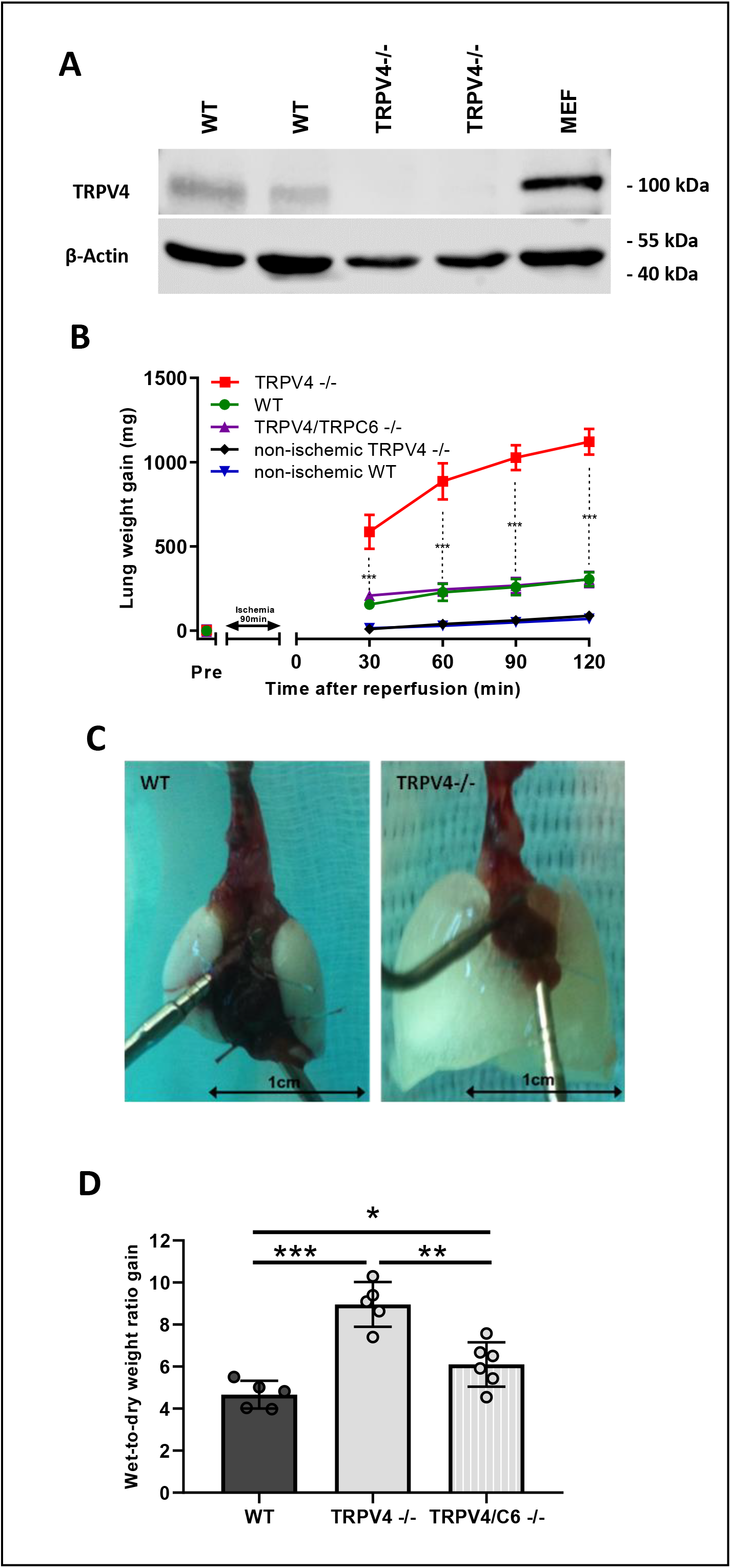
Ablation of TRPV4 increases ischemia-induced edema formation in mouse lungs. (**A**) TRPV4 protein expression in mouse lungs was evaluated by immunoblotting in whole lung lysates of wild-type (WT) and TRPV4-deficient (TRPV4−/−) mice using a TRPV4-specific antiserum. Murine embryonic fibroblasts (MEF) cells served as additional positive control. Expression of β-actin was used as loading control. (**B**) Constant weight measurement of ischemic and non-ischemic WT (n = 5) and TRPV4−/− (n = 5) and TRPV4/TRPC6-double deficient (TRPV4/C6−/−, n = 6) isolated perfused lungs. (**C**) Representative images of WT and TRPV4−/− lungs after ischemia. (**D**) Wet-to-dry weight ratio gains of TRPV4−/− and TRPV4/TRPC6−/− lungs compared to WT controls. Data represent means ± SEM of at least 5 lungs for each genotype. Significance between means was analyzed using two way ANOVA (**B**) or two tailed unpaired Student’s t-test (**D**) and indicated as *** for p<0.001, ** for p<0.01 and * for p<0.05.

### TRPV4 is expressed in alveolar epithelial type I (ATI) and type II (ATII) cells

As TRPV4 is highly expressed in lung endothelium, and its activation results in an increase of endothelial permeability (reviewed in (*Simmons et al., 2018*)), we focused on its possible function in epithelial cells representing the second natural barrier regulating edema formation. Analysis of mice carrying an eGFP-reporter protein under the control of the TRPV4 promoter/enhancer region revealed expression of TRPV4 protein in endothelium, bronchial as well as alveolar epithelium (*Figure 2A*). In the bronchial epithelium we detected TRPV4 in ciliated cells by co-staining with a β-tubulin IV antibody (*Figure 2-figure supplement 1A-C*). Club cells as well as neuroendocrine cells did not show TRPV4 expression (*Figure 2-figure supplement 1D-I*). In the alveoli, co-staining experiments with an antibody directed against aquaporin-5 (AQP5) (*Figure 2B*) - a marker protein of ATI cells involved in lung septa formation *(Dobbs et al., 2010*) - revealed a red staining indicative of AQP-5 expression in the plasma membrane and an additional green staining of the cytosol reflecting TRPV4 expression in these cells. Moreover, direct quantification of TRPV4 mRNA revealed similar expression levels in ATII cells as in lung endothelial cells (EC), but lower mRNA expression in pulmonary murine lung fibroblasts (pmLF), and precapillary arterial smooth muscle cells (PASMC) (*Figure 2C*).

**Figure 2.**
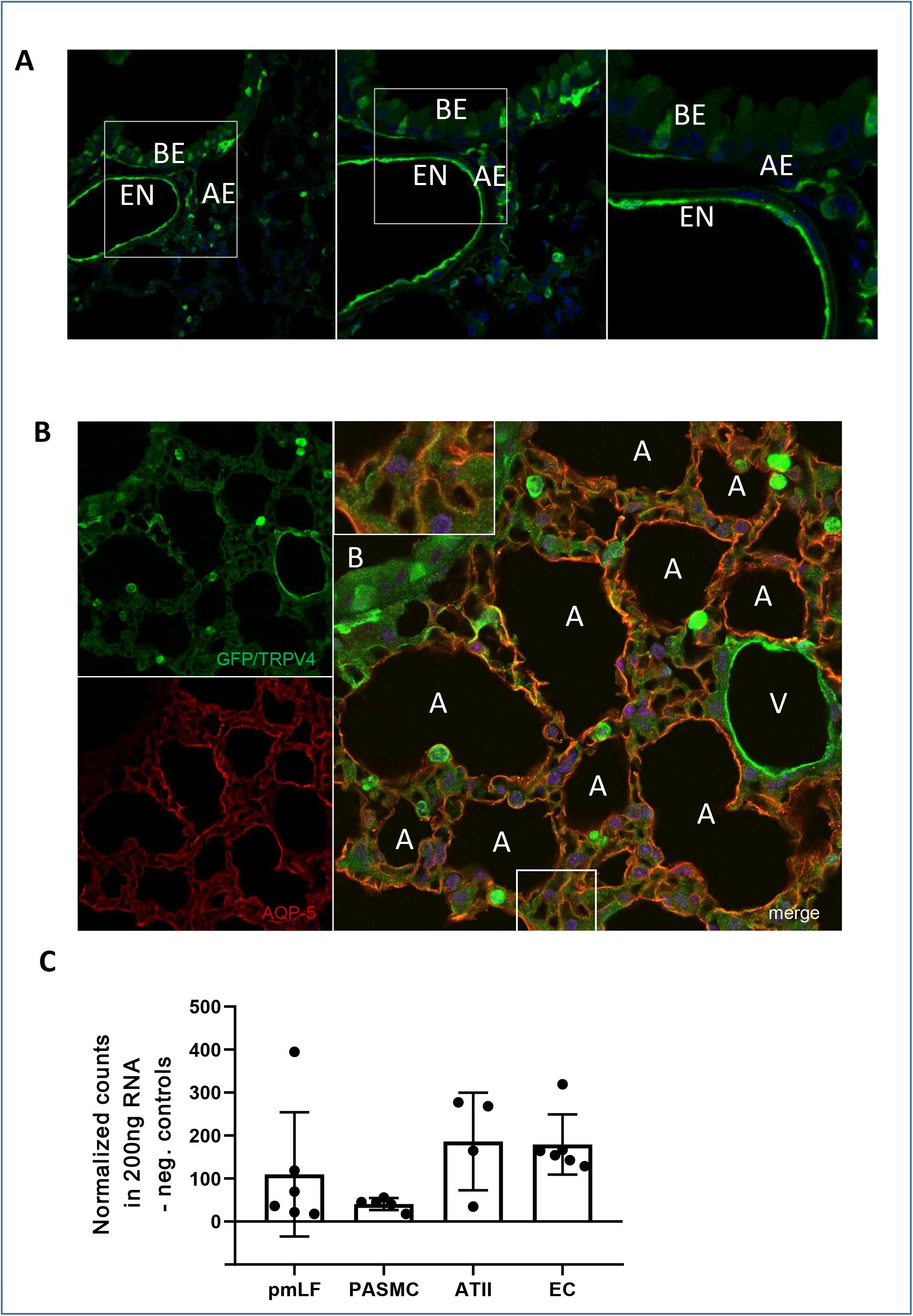
TRPV4 and aquaporin-5 (AQP-5) expression in mouse lungs. (**A**) GFP staining (green) by fluorescent-coupled GFP-specific antibodies in lung cryosections of TRPV4eGFP reporter mice reveals expression of TRPV4 in the cytoplasm of cells in lung endothelium (labelled EN) as well as bronchial (BE) and alveolar epithelium (AE). Nuclei staining was performed by Hoechst-dye (blue). Magnifications of 24, 50 and 100 x are shown. (**B**) Lung cryosections from TRPV4eGFP-reporter mice were stained with fluorescent-coupled antisera directed against GFP and AQP-5. Confocal images were obtained after excitation at 488 nm (for eGFP, left upper panel marked as green staining) or after excitation at 561 nm (for AQP-5, left lower panel marked as red staining). Both images at a magnification of 40 x were merged (right panel). Nuclei staining was performed by Hoechst-dye (blue). A, alveolus; B, bronchus; V, vasculature. Inset is showing the lower boxed region in a higher magnification (80 x) **(C)** TRPV4-mRNA quantification in lung cells by the nanostring®-technology. ATII, aleveolar type II cells; EC, endothelial cells; PASMC, precapillary arterial smooth muscle cells; pmLF, primary murine lung fibroblasts. Data represent means ± SEM from at least 3 independent cell isolations.

### Loss of TRPV4 resulted in decreased aquaporin-5 expression in TRPV4−/− lungs

Staining of lung slices with fluorescence-coupled antibodies specific for the water conducting channel AQP-5 revealed lower total expression levels in septa-forming ATI cells and reduced plasma membrane localization in TRPV4−/− lungs compared to WT lungs (*Figure 3A-E*). These results were confirmed by Western blotting analysis of lung lysates probed with an AQP-5-specific antibody (*Figure 3F-G*). In clear contrast to these results, protein levels of aquaporin-1 (AQP-1) - the major aquaporin channel in the microvascular endothelium - were not significant different in TRPV4−/− endothelial compared to WT cells (*Figure 3-figure supplement 1A-E*).

**Figure 3.**
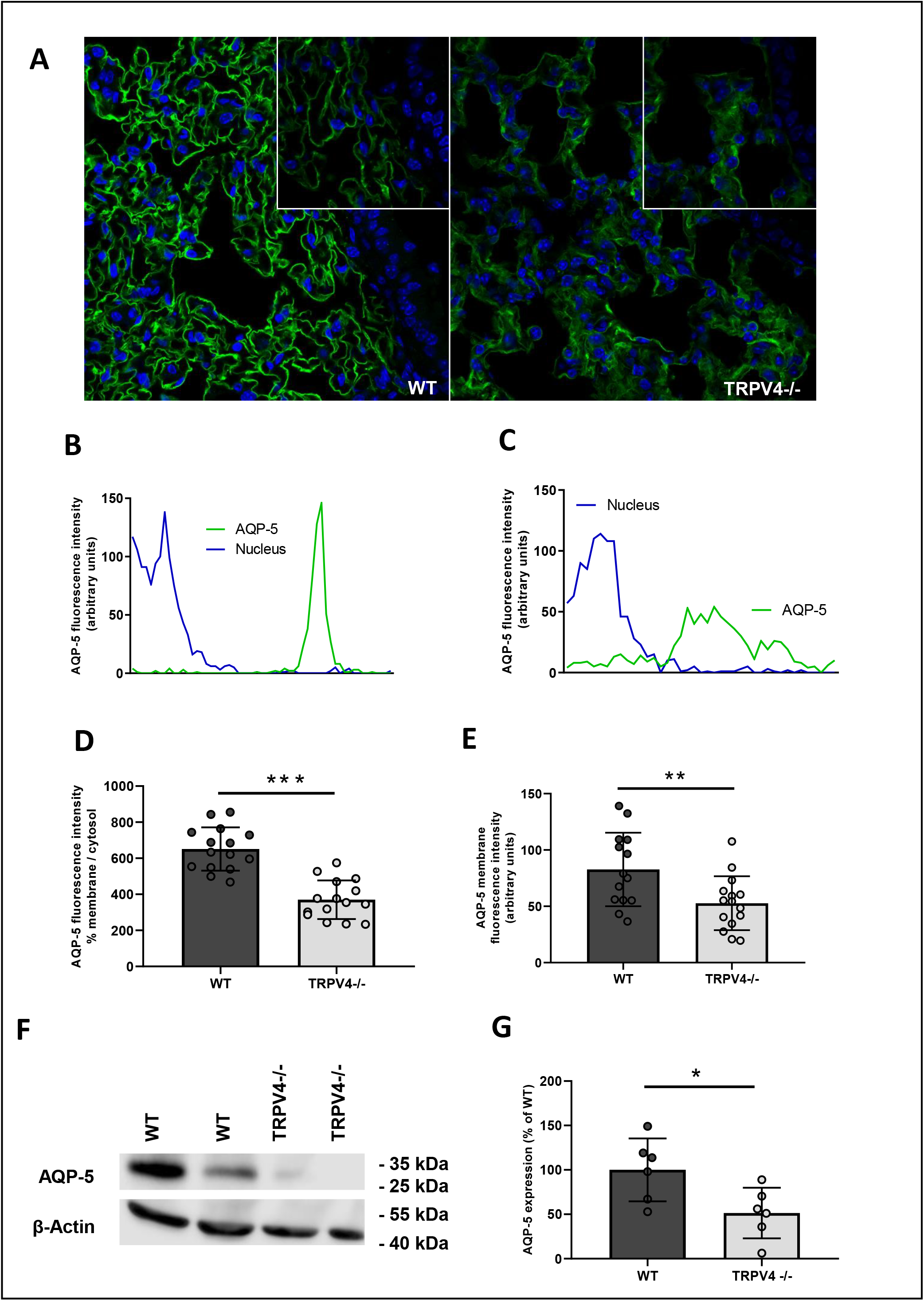
Aquaporin-5 (AQP-5) expression and translocation to the plasma membrane in WT and TRPV4−/− alveolar epithelial type I (ATI) cells. (**A**) Cryosections of WT and TRPV4−/− lungs stained with an AQP-5 specific fluorescent-coupled antibody. Nuclei staining was performed by Hoechst-dye (blue). Magnification of 40 x (large images) and 100 x (small images) are shown. Representative histograms used for quantification of AQP-5 protein in the plasma membrane of WT (**B**) and TRPV4-deficient ATI cells (**C**). Summaries of AQP-5 protein expression in plasma membranes in relation to the cytosol (AQP-5 fluorescence intensity % membrane/cytosol (**D**)) and total expression in plasma membranes (**E**). Representative Western blot analysis of AQP-5 expression in WT and TRPV4−/− whole lung lysates (**F**) and quantification of AQP-5 (**G**). Data represent means ± SEM from at least 6 lungs for each genotype. Significance between means was analyzed using two tailed unpaired Student’s t-test and indicated as *** for p<0.001, ** for p<0.01 and * for p<0.05.

### Identification of currents induced by the TRPV4 activator GSK1016790A only in primary ATII cells from WT mice producing more pro surfactant protein-C (pSP-C) than TRPV4−/− cells

To investigate the role of TRPV4 on a cellular level, we first isolated ATII epithelial cells (*Figure 4A*) from WT and TRPV4−/− mice, which showed no morphological differences. ATII cells were identified by staining with fluorophore-coupled antibodies directed against pro surfactant protein-C (pSP-C) (*Figure 4B*), which is produced by ATII cells (reviewed in (*Fehrenbach, 2001*)). Patch clamp analysis of primary ATII cells revealed significantly larger currents induced by the selective TRPV4 activator GSK1016790A (reviewed in (*Dietrich, 2019*)) only in WT cells. Moreover, GSK1016790A-induced currents in TRPV4−/− cells were not significantly from the basal currents in WT cells (*Figure 4C-D*). Western blotting analysis of protein lysates from ATII cells revealed lower pSP-C levels in TRPV4−/− ATII cells compared to WT cells (*Figure 4E-F*). We then differentiated ATII to ATI cells by growing them to confluency in plastic cell culture dishes for at least 6 days as described (*Mutze et al., 2015*) (*Figure 4G*). After 6 days WT cells expressed AQP-5 protein as an ATI cell marker (*Figure 4H*).

**Figure 4.**
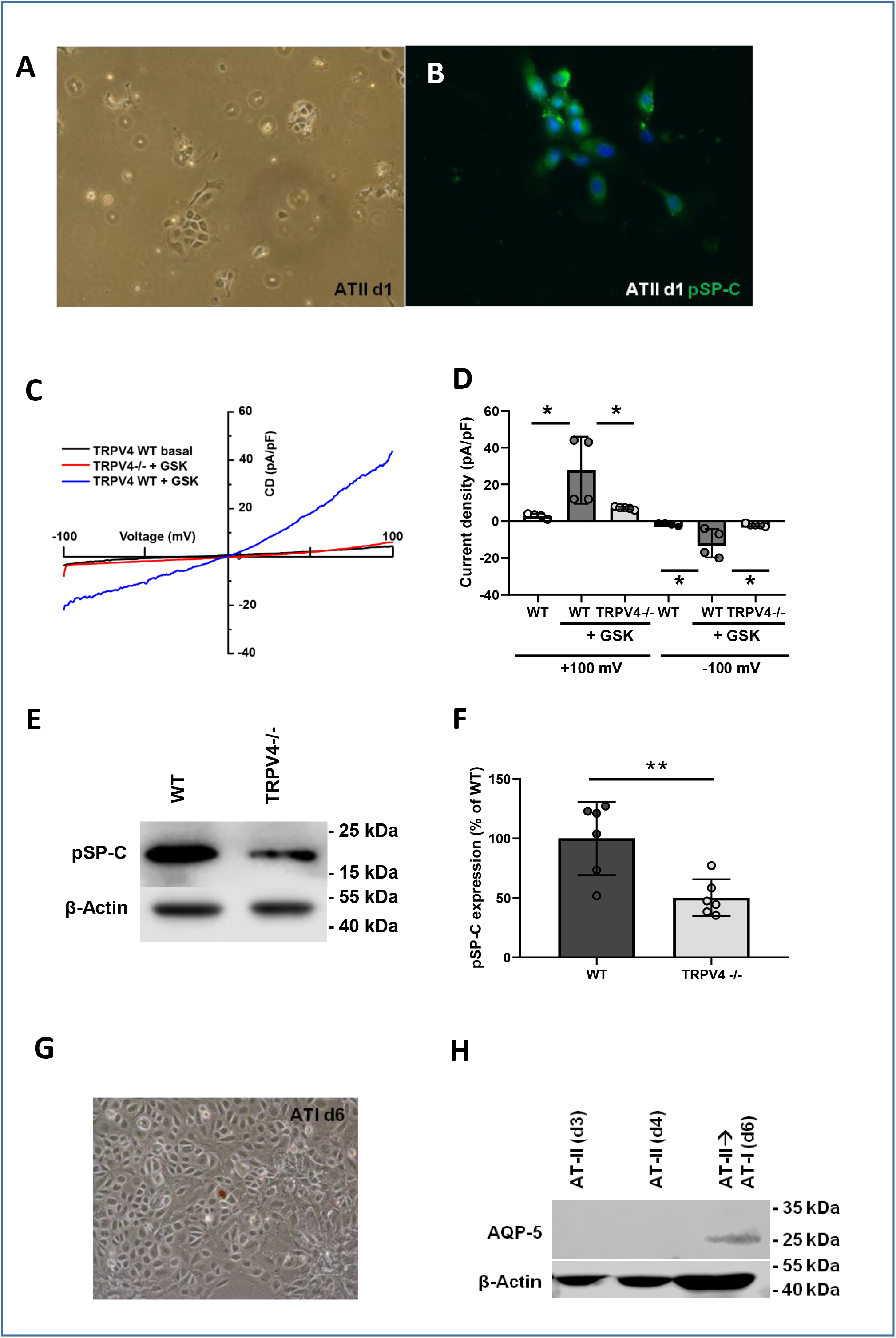
Identification of alveolar type II (ATII) cells and differentiation to alveolar type I (ATI) cells. **(A**) Representative cell cluster one day after isolation in a phase contrast image and stained with a fluorescent-coupled specific pro-surfactant protein-C (pSP-C) antibody (**B**). Nuclei staining was performed by Hoechst-dye (blue). Electrophysiological whole-cell measurements of basal and GSK1016790A-induced current densities in WT and TRPV4−/− primary ATII cells (**C-D**). Representative current density–voltage curves of wild-type (grey, blue traces) and TRPV4−/− (red trace) ATII cells before (grey trace) and during application of 1 mM GSK1016790A (GSK, blue and red traces) (**C**). Summary of current densities at +/-100 mV before (white bars) and after application of GSK1016790A analyzed in WT (black bars) and TRPV4−/− (grey bars) ATII cells (**D**). Representative western blot analysis of pSP-C expression in WT and TRPV4−/− ATII cells (**E**) and summary of pSP-C expression in TRPV4−/− and WT ATII cells (**F**). Image of confluent cells on day 6 after ATII cell isolation (**G**) and analysis of AQP-5 expression in cells grown for 3, 4 and 6 days in plastic cell culture dishes by Western-blotting (**H**). Expression of β-actin was used as loading control in each blot. Data represent means ± SEM from at least 3 independent cell preparations of 5 mice each. Significance between means was analyzed using one way ANOVA (**C**) or two tailed unpaired Student’s t-test (**F**) and indicated as ** for p<0.01 and * for p<0.05.

### TRPV4−/− ATI cells produce less AQP-5, show reduced nuclear localization of nuclear factor of activated T-cells (NFAT) as well as decreased cell migration and adhesion

As already shown in lung sections of TRPV4−/− mice AQP-5 expression was reduced in TRPV4−/− cells (*Figure 4A-B*). To test if TRPV4−/− ATII cells are able to differentiate to ATI cells, we analyzed the expression of podoplanin (T1α), another ATI cell marker protein. Notably, podoplanin expression was not significantly different in TRPV4−/− ATII cells differentiated to ATI cells (*Figure 4C-D*).To further analyze ATI cell function, we quantified nuclear NFATc1 levels, which were significantly reduced in TRPV4−/− cells (*Figure 5E-F*). Moreover, cell migration analyzed by gap closure in in-vitro experiments was clearly slowed down in TRPV4-deficient ATI cells compared to WT cells (*Figure 5G-H*). As an additional line of evidence we transfected ATII cells with TRPV4-specific or control siRNAs, differentiated them to ATI cells and quantified cell migration in the same way (*Figure 5-figure supplement 1*). Most interestingly, we obtained similar results as cells transfected with TRPV4-siRNA showed a significantly slowed migration compared to non-transfected cells as well as cells transfected with the control siRNAs. Moreover, as determined by electrical cell impedance sensing (ECIS), sub-confluent ATI cells also showed reduced cell barrier function (*Figure 5 I*).

**Figure 5.**
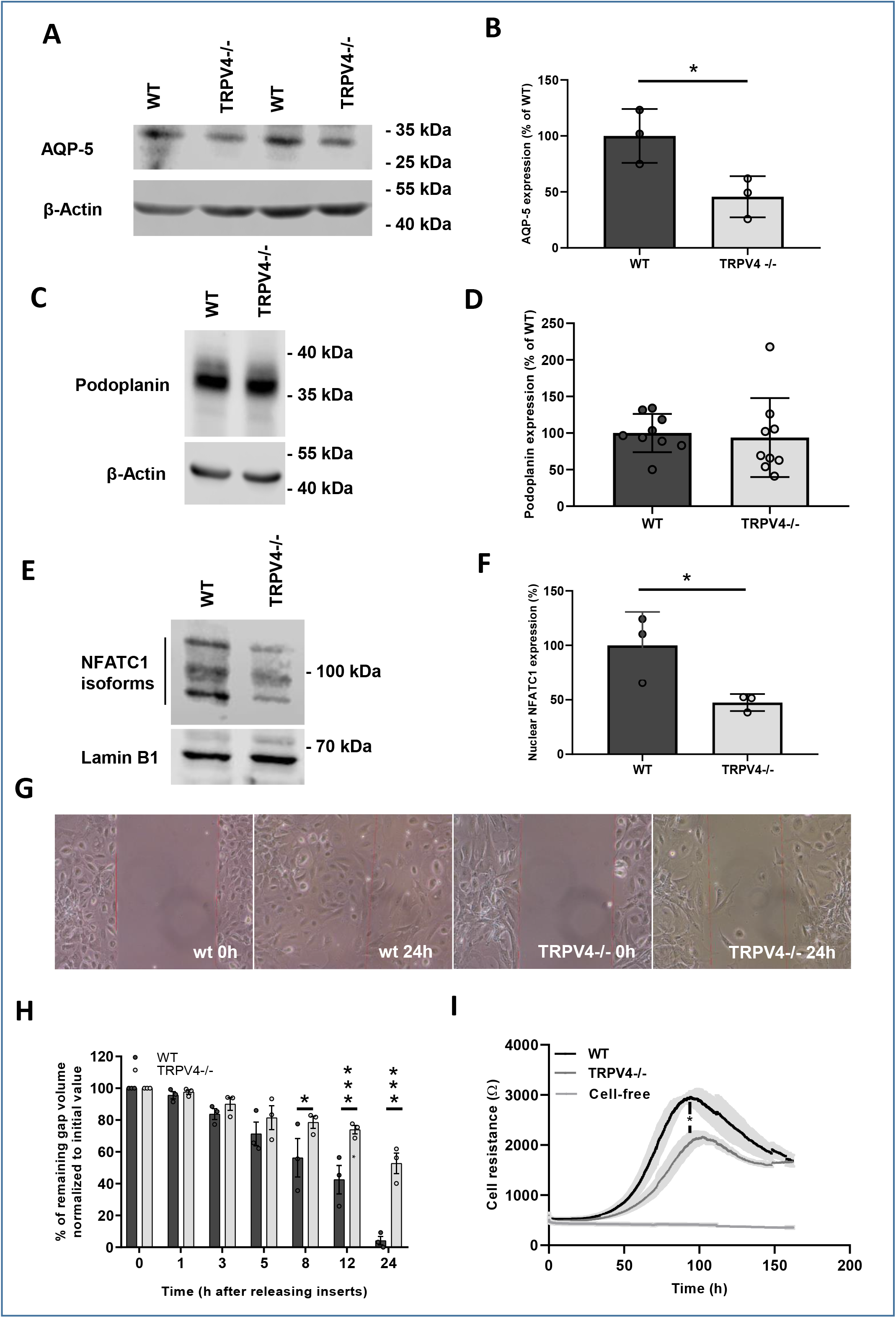
Nuclear localization of nuclear factor of activated T-cells (NFAT) in and migration and adhesion of TRPV4-deficient and WT ATI cells. Representative western blot analysis of AQP-5 expression in WT and TRPV4−/− ATII cells differentiated to ATI cells (**A**) and summary of AQP-5 expression in these cells (**B**). Representative western blot analysis of podoplanin expression – another ATI cell marker - in WT and TRPV4−/− ATII cells differentiated to ATI cells (**C**) and summary of podoplanin expression in these cells (**D**). Representative western blot analysis of nuclear NFATc1 localization in WT and TRPV4−/− ATI cells (**E**) and summary of nuclear NFAT localization in these cells (**F**). Lamin B1 expression served as loading control. Representative images of a migration assay after removing the insert at time pints 0 and 24 h (**G**). Summary of remaining gap values normalized to initial values quantified in migration assays of TRPV4−/− and WT ATI cells after removing inserts at 0, 1, 3, 5, 8, 12 and 24 h (**H**). Electrical cell resistance was quantified with an ECIS device for WT and TRPV4−/− ATI cells (cell preparation from 4 mice) for 160 h (**I**). Data represent means ± SEM from at least 3 independent cell preparations of 5 mice each. Significance between means was analyzed using two tailed unpaired Student’s t-test and indicated as *** for p<0.001 and * for p<0.05.

## TRPV4−/− mice expose emphysema-like lung structures and altered lung function

To analyze differences in lung anatomy as a consequence of altered ATI cell function, we quantified mean chord lengths in histological lung sections (*Figure 6A*). TRPV4-ablation significantly increased mean chord length of the alveolar lumen in adult (three to twelve months old (*Figure 6C-D*)) mice compared to WT lungs, while young mice (three weeks old) showed no differences (*Figure 6B*). Moreover, lung function was altered (*Figure 6E-H*): TRPV4−/− lungs showed increased inspiratory capacity and compliance (*Figure 6E, G, H*) as well as decreased elastance (*Figure 6F*) significantly different from WT mice of the same age.

**Figure 6.**
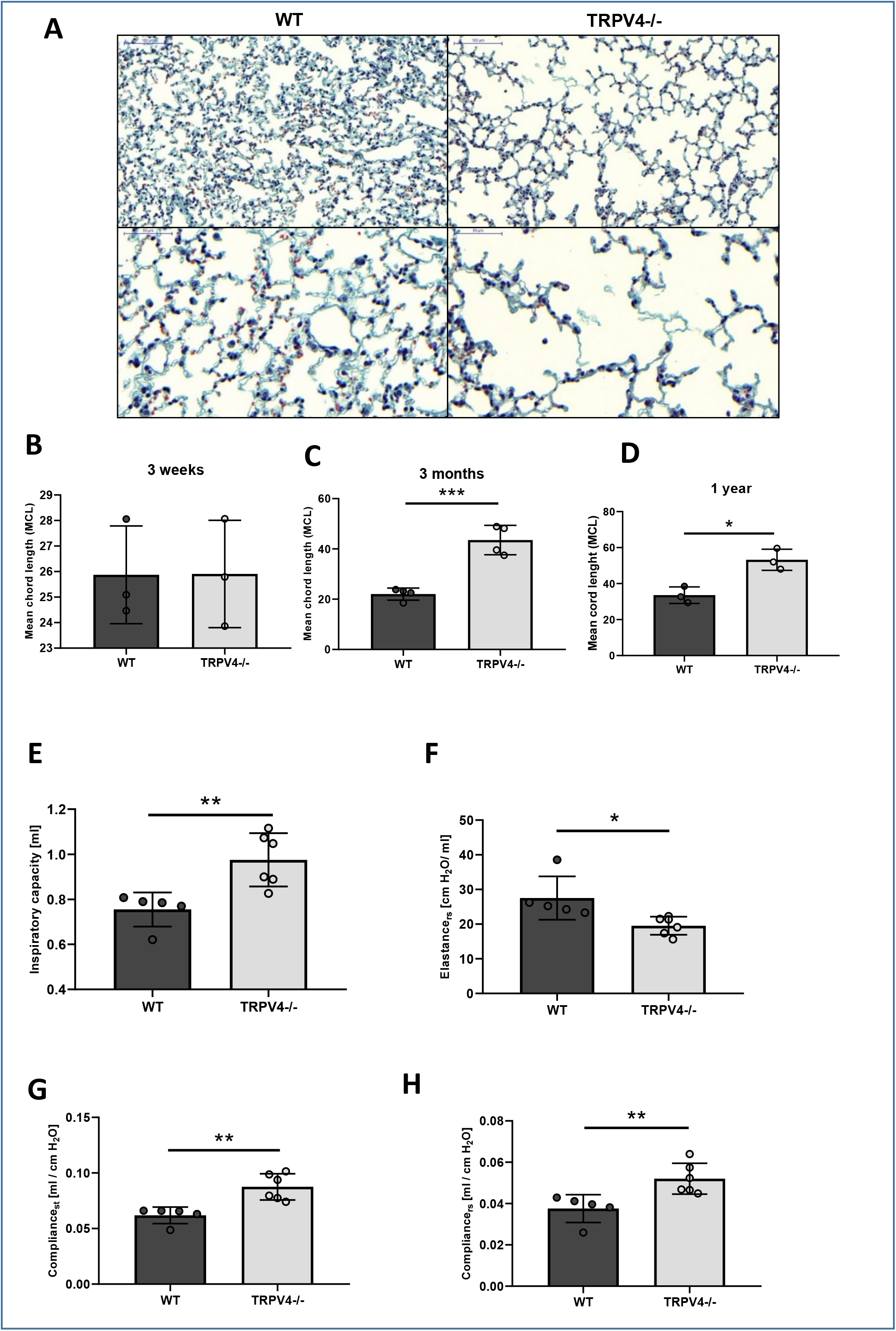
Chord lengths and lung function of WT and TRPV4−/− mice. Representative images of masson-trichrome-stained lung sections from 3 months old WT and TRPV4−/− mice at 20 x magnification (upper panels) and at 40 x magnification (lower panels) (**A**). Quantification of mean chord lengths of 3 weeks (**B**), 3 months (**C**) and 1 year old (**D**) WT and TRPV4−/− mice. Inspiratory capacity (IC (**E**)), elastance of the respiratory system (E_rs_ (**F**)), static compliance (C_st_ (**G**)) and compliance of the respiratory system (C_rs_ (**H**)) of 6 months old WT and TRPV4−/− mice. Data represent means ± SEM from at least 3 mice. Significance between means was analyzed using two tailed unpaired Student’s t-test and indicated as *** for p<0.001, ** for p<0.01 and * for p<0.05.

## Discussion

Although TRPV4 is highly expressed in lungs, its exact function is still elusive (reviewed in (*Dietrich, 2019*)). Activation of TRPV4 in endothelial cells by mechanical stress e.g. stretching (*Hamanaka et al., 2007; Michalick et al., 2017; Yin et al., 2008*) as well as oxidative stress by exposure to H_2_O_2_ (*Suresh et al., 2015*) resulted in an increased Ca^2+^ influx mediated by the channel and an increase in endothelial permeability conducive to lung edema (reviewed in (*Simmons et al., 2018*)). Along these lines, pharmacological blockade of TRPV4, e.g. by the specific blocker HC-067047, decreased endothelial rises in intracellular Ca^2+^ and protected mice from vascular leakage and lung injury (*Michalick et al., 2017*). Here, we quantified ischemia-reperfusion (IR)-induced edema as one of the most common and significant causes of morbidity and mortality after lung transplantation (*de Perrot et al., 2003*) using the isolated perfused mouse lung (IPL) (*Weissmann et al., 2012*). Much to our surprise, TRPV4−/− lungs were not protected from IR-induced lung edema as observed in TRPC6−/− mice (*Weissmann et al., 2012*). On the contrary, genetic TRPV4 ablation resulted in a robust increase in lung edema (*Figure 1B*) and a higher wet-to-dry weight ratio gain (*Figure 1D*) when compared to control wild-type (WT) mice. Barrier function was rescued by consecutive breeding TRPV4−/− mice with TRPC6−/− mice, because lung edema formation in double deficient mice was similar to WT animals (*Figure 1B*).

As TRPV4 activation in endothelial cells has been shown to result in higher edema formation (*Simmons et al., 2018*), we focused on the lung epithelium, another physiological cell barrier in the lung. Recent publications indicate an epithelial function of the channel opposed to that in endothelium i.e. stabilization of the epithelial barrier in the skin (*Akazawa et al., 2013*), the urogenital tract (*Janssen et al., 2016*) and the corneal epithelium (*Martinez-Rendon et al., 2017*). We demonstrated TRPV4 expression in alveolar epithelial type I (ATI) and type II (ATII) cells (*Figure 2B-C*). Our further molecular analysis corroborated a functional link between TRPV4 and AQP-5, a water conducting channel expressed in ATI cells (*Dobbs et al., 1998*). Hypotonic solutions increased the association and surface localization of TRPV4 and AQP-5 in salivary gland cells (*Liu et al., 2006*) and AQP-5 expression is regulated by TRPV4 in lung epithelial cells (*Sidhaye et al., 2006*). Most interestingly, the expression and plasma membrane translocation of AQP-5 channels in ATI cells were significantly reduced (*Figure 3A-G*). To analyze TRPV4 function on a cellular level, we isolated ATII cells identified by their expression of pro surfactant protein-C (pSP-C) (*Figure 4A-B*). We were able to identify significantly larger currents induced by the TRPV4 activator GSK1016790A in WT but not in TRPV4−/− ATII cells (*Figure 4C-D*). To our knowledge these data show for the first time that TRPV4 channels are not only expressed, but are also functional in ATII cells. Quantifying pSP-C levels by Western Blotting revealed a reduced expression in TRPV4−/− cells compared to WT cells (*Figure 4E-F*). The role of surfactant proteins in the prevention of alveolar edema by reducing surface tension as a driving force for fluid flow across the air blood barrier, is still a matter of debate (*Hills, 1999*), but might also explain exaggerated edema formation in TRPV4−/− mice.

Next, we differentiated ATII to ATI cells (*Mutze et al., 2015*), monitored by the expression of two ATI cell markers: AQP-5 and podoplanin. As AQP-5 protein expression was reduced in TRPV4−/− ATI cells (*Figure 5A-B*) while podoplanin levels were not altered (*Figure 5C-D*), it seems rather unlikely that TRPV4-deficency and/or a reduction of pSP-C expression results in reduced ATII to ATI differentiation in general. Plasma membrane translocation of AQP-5 as well as AQP-5 expression may depend on nuclear localization of the transcription factor nuclear factor of activated T cells (NFAT) by a rise of intracellular Ca^2+^ via TRPV4 similar to TRPC channels (*Curcic et al., 2019*). Therefore, we quantified nuclear NFAT levels and detected significantly lower levels in TRPV4−/− cells in comparison to WT control cells (*Figure 5E-F*). A major breakthrough in our understanding of AQP-5 function for water transport across apical membranes of ATI cells, was the analysis of AQP-5-deficient mice (*Ma et al., 2000*). Although lack of AQP-5 entailed a 10-fold decrease in alveolar permeability in response to an osmotic gradient, AQP-5−/− mice are indistinguishable from WT mice with regard to hydrostatic pulmonary edema as well as isosmolar fluid transport from the alveolar space (*King et al., 2000; Ma et al., 2000*). Cognizant of this scenario, a role for AQP-5 in the clearance of fluid from the alveolar space after IR-induced lung edema cannot entirely be ruled out, but appears to be unlikely, and we tried to dissect other additional mechanisms for the vulnerability of TRPV4−/− lungs to edema formation.

As two reports demonstrated decreased migration of human epithelial ovarian cancer (*Yan et al., 2014*) or endometrial adenocarcinoma cells (*Jiang et al., 2012*) after downregulation of AQP-5, we set out to quantify cell migration of ATII cells differentiated to ATI cells. TRPV4−/− ATI cells showed a clear deficit in closing gaps by cell migration after releasing inserts compared to WT cells (*Figure 5G-H*). In additional experiments we were able to reproduce these results in cells transfected with TRPV4-siRNAs compared to non-transfected cells as well as cells transfected with control siRNAs (Supplementary Fig. 3). These data clearly identify acute TRPV4 ablation as the initial cause of reduced cell migration. Moreover, cell resistance as analyzed by electrical cell impedance sensing (ECIS) was significantly reduced in growing TRPV4−/− ATI cells in contrast to WT cells (*Figure 5I*), while no differences in morphology and proliferation rates were detected.

ATII cells are able to differentiate to ATI cells after lung injury during repair processes in adult mice (*Desai et al., 2014*) to reestablish barrier function of the lung alveolus. Thus, we analyzed lung alveolar histology in WT and TRPV4−/− lungs in young and adult mice. Mean chord length as a measure of alveolar size was increased in adult (3 months – 1 year old) but not in young (3 weeks old) TRPV4−/− mice compared to WT mice of the same age (*Figure 6A-D*). We concluded that differences were not caused by defects in embryonic lung development, but were due to ongoing growth and repair processes in adult animals. Most interestingly, the emphysema-like changes in lung morphology were also detected in SP-C-deficient mice (*Glasser et al., 2003*), raising the possibility that reduced pSP-C levels in TRPV4−/− ATII cells may also contribute to the phenotype. In the same vein, adult TRPV4−/− mice showed altered lung function with increased inspiratory capacity and compliance as well as decreased elastance (*Figure 6E-H*) compared to WT mice of the same age. Loss of septa formation because of reduced pSP-C levels in adult TRPV4−/− mice may be responsible for decreased clearance of fluid from the alveolar space and may therefore explain higher edema formation in TRPV4−/− lungs.

In summary, loss of TRPV4 channels in alveolar epithelial cells results in decreased production of pSP-C production in ATII and lower AQP-5 expression and membrane localization in ATI cells. The latter proteins are likely to be involved in continuously ongoing repair processes in adult mice, resulting in emphysema-like changes in TRPV4−/− mice. These molecular events may define a protective function of TRPV4 channels against lung edema formation, in clear contrast to their detrimental role in endothelial cells.

## Materials and Methods

### Animals

TRPC6−/−, TRPV4−/− (B6.199×1-Trpv4^tm1MSZ^ from Riken BioResource Center (RBRC01939) (Mizuno et al., 2003; Suzuki et al., 2003)), TRPC6/TRPV4−/− and TRPV4eGFP reporter mice (Tg(TRPV4-EGFP)MT43Gsat/ Mmucd from MMRC) were bred as previously described (*Dietrich et al., 2005; Gong et al., 2003; Suzuki et al., 2003*). All animal experiments were approved by the governmental authorities.

### Isolated, perfused mouse lung (IPL)

Quantification of edema formation in isolated perfused mouse lungs (IPL) were done as described (*Weissmann et al., 2012*). In brief, mice were anesthetized by intraperitoneal injection of ketamine (100 mg/kg body weight (bw)), xylazine (0,7 mg/kg bw) and anticoagulated with heparin (500 iU./kg bw). Animals were intubated via a small incision in the trachea, ligated and ventilated with room air using the VCM type 681 (positive end−expiratory pressure, 3 cmH_2_O; positive end-inspiratory pressure 3 cmH_2_O; respiratory rate was 90 breaths/min). The sternum was opened, the ribs were spread and the right ventricle was incised to place the air-free perfusion catheter into the pulmonary artery. After ligation the perfusion was started with 0.5 ml/min perfusion solution (7.19 g sodium chloride, 0.33 g potassium chloride, 0.27 g magnesium hexahydrate, 0.36 g calcium chloride dihydrate, 0.15 g potassium dihydrogen orthophosphate, 2.67 g glucose monohydrate, 51.28 g hydroxyethyl starch 200000/05 ad 1000 ml with aqua ad injectabilia, use 0.1848 mg/ml sodium hydrogen carbonate to adjust pH to 7.3) using an ISAMATEC Tubing Pump. A second perfusion catheter was introduced in the left ventricle and secured by ligation. The lung, together with the trachea and the heart were excised from the thorax in one piece and transferred to a 37 °C temperature-equilibrated housing chamber of the perfused mouse lung model (IPL-2, Hugo Sachs Elektronik/Harvard Apparatus (March-Hugstetten, Germany)). The perfusion was slowly raised stepwise to 2 ml/min and monitored with the PLUGSYS® TAM-A/P75 type 17111. Weight changes were constantly measured with the edema Balance Module/ EBM type 713. Data was monitored with the Pulmodyn software.

### Analysis of functional parameters of the respiratory tract

Mice were anesthetized with ketamine (270 mg/kg bw) and xylazin (11 mg/kg bw), intratracheally intubated through a small incision of the trachea and connected to the flexiVent system (Scireq, Montreal, Canada).

### Immunohistochemistry

Mouse lungs were inflated with 4% (m/v) paraformaldehyde in PBS and processed for paraffin or O.C.T compound (Tissue-Tek, Sakura finetek, Torrance, CA, USA) embedding. Paraffin-embedded tissue sections (3 μm) were cut using a microtome (Zeiss, Göttingen, Germany), mounted on glass slides, deparaffinized in xylene and rehydrated in graded alcohol. Masson Goldner trichrome staining (Masson Goldner Trichrome Staining Kit, Carl Roth 3459) was done as described in the manufacture’s instruction manual with iron hematoxylin solution for 8 min, Goldner’s stain 1 for 6 min, Goldner’s stain 2 for 1 min and Goldner’s stain 3 for 5 min. After dehydration in 100% EtOH and clearing in xylol twice for 1 min, the sections were mounted using the Roti-Histokit II (Carl Roth T160.2). Sections were analyzed by design-based stereology using an Olympus BX51 light microscope equipped with the new Computer Assisted Stereological Toolbox (newCAST, Visiopharm) as described (John-Schuster et al., 2014). For mean chord length (MCL) measurements, 10-20 frames were selected randomly across multiple sections by the software, using the 20x objective, and superimposed by a line grid and points. The intercepts of lines on alveolar wall (Lsepta) and points localized on air space (Pair) were counted and calculated as MCL (∑Pair × L(p) / ∑Isepta × 0.5, where L(p) is the line length per point). Cryo-embedded lungs were cut in 10 µm sections on a cryostat (Leica, Wetzlar, Germany), mounted on glass slides and surrounded with a hydrophobic pen (Vector Laboratories, California, USA). After washing with PBS the sections were blocked for 30 min. in PBS containing 0,2% Triton X-100 and 5% NGS. Incubation with primary antibody was done at 4 °C overnight and secondary antibody at room temperature (RT) for 1 h. Antibodies were diluted in blocking solution. After nuclei staining with Hoechst-dye (2 µg/ml) for 5 min at RT followed by sufficient washing the sections were mounted in Roti-Histokit II. Used antibodies and dilutions were: anti-GFP (chicken, Thermo Fisher, A10262, 1:200), anti-β-tubulin IV (rabbit monoclonal, Abcam, 179509) 1:1600), anti-aquaporin-1 (rabbit, Alomone Labs, AQP-001, 1:100), anti-aquaporin-5 (rabbit, Alomone Labs, AQP-005, 1:100), anti-CC10 (mouse, Santa Cruz, E-11, 1:200) anti-chicken (goat, Thermo Fisher, A11039, 1:400), anti-CGRP (goat, 1:400, Acris, BP022), anti-pro-surfactant-protein-C (rabbit, 1:20,000, Merck, AB3786, pSP-C), anti-rabbit IgG (goat, coupled to Alexa 488, Thermo Fisher, A32731, 1:500 and donkey, coupled to Cy3, Merck Millipore, AP182C, 1:1000), anti-goat IgG (donkey, Life Technologies, A11058, 1:400). ^For direct labeling of the anti-CC10 antibody Zenon Alexa Fluor 546 mouse IgG^1 ^kit^ was used according to the manufacturer’s recommendations (Invitrogen, 25004). Stained cryosections were analyzed on an epifluorescence microscope (Zeiss Imager.M2, Carl Zeiss, Jena, Germany) and on a confocal microscope (LSM 880, Carl Zeiss). For membrane localization analysis staining intensity was analyzed along a line from the nucleus into the cytosol and the plasma-membrane.

### Primary murine alveolar epithelial cells

Isolation of alveolar epithelial cells type 2 (ATII) was done as described (Corti et al., 1996; Dobbs, 1990; Mutze et al., 2015). In brief, lungs were flushed via a catheter through the pulmonary artery with 0.9% NaCl solution (B. Braun Melsungen AG, Melsungen, Germany), inflated with 1 ml dispase (BD Bioscience, San Jose, CA) followed by 500 µl 1% low-melting-point agarose (Sigma-Aldrich, St Louis, MO) and incubated for 1 h at RT. Subsequently, lung lobes were separated and dissected using two forceps, filtered through 100 µm, 20 µm and 10 µm nylon filters (Sefar, Heiden, Switzerland) and centrifuged for 10 min at 200 x g. Cell pellets were resuspended in DMEM (Sigma-Aldrich) and plated on CD45 and CD16/32 (BD Bioscience) coated culture dishes for a negative selection of macrophages and lymphocytes and incubated for 30 min at 37 °C. Non-adherent cells were collected and seeded on uncoated dishes to negatively select fibroblasts at 37 °C for 25 min. Cells were collected and live cells were counted by trypan blue staining in a Neubauer counting chamber. Two x 10^6^ cells/well of a 6-well plate were seeded in DMEM containing 10% FCS (Invitrogen, Carlsbad; USA), 1% HEPES (Carl Roth, Karlsruhe, Germany) and 1% penicillin/streptomycin (Lonza, Basel, Switzerland) and used for analysis or grown for at least 6 d for ATI cell differentiation. ATII cells were transfected with 1 μM Accell SMARTpool siRNA for TRPV4 (in starving medium, 0.1% FCS) two days after isolation. On day 6, the cells were washed once and kept in starving medium. A non-coding pool of the Accell siRNA in starving medium served as control (see *Table 1* for siRNA sequences).

**Table 1.**
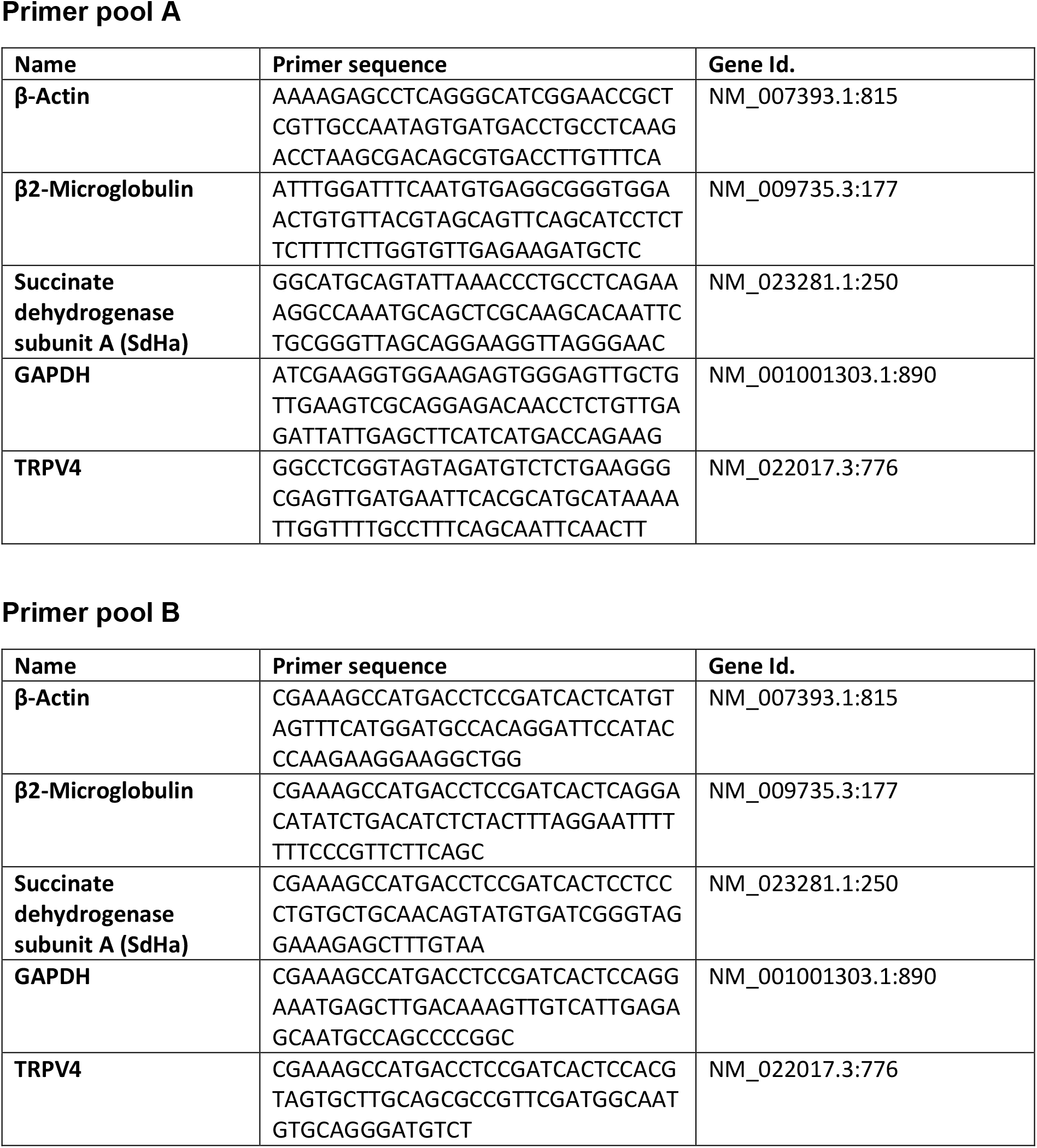
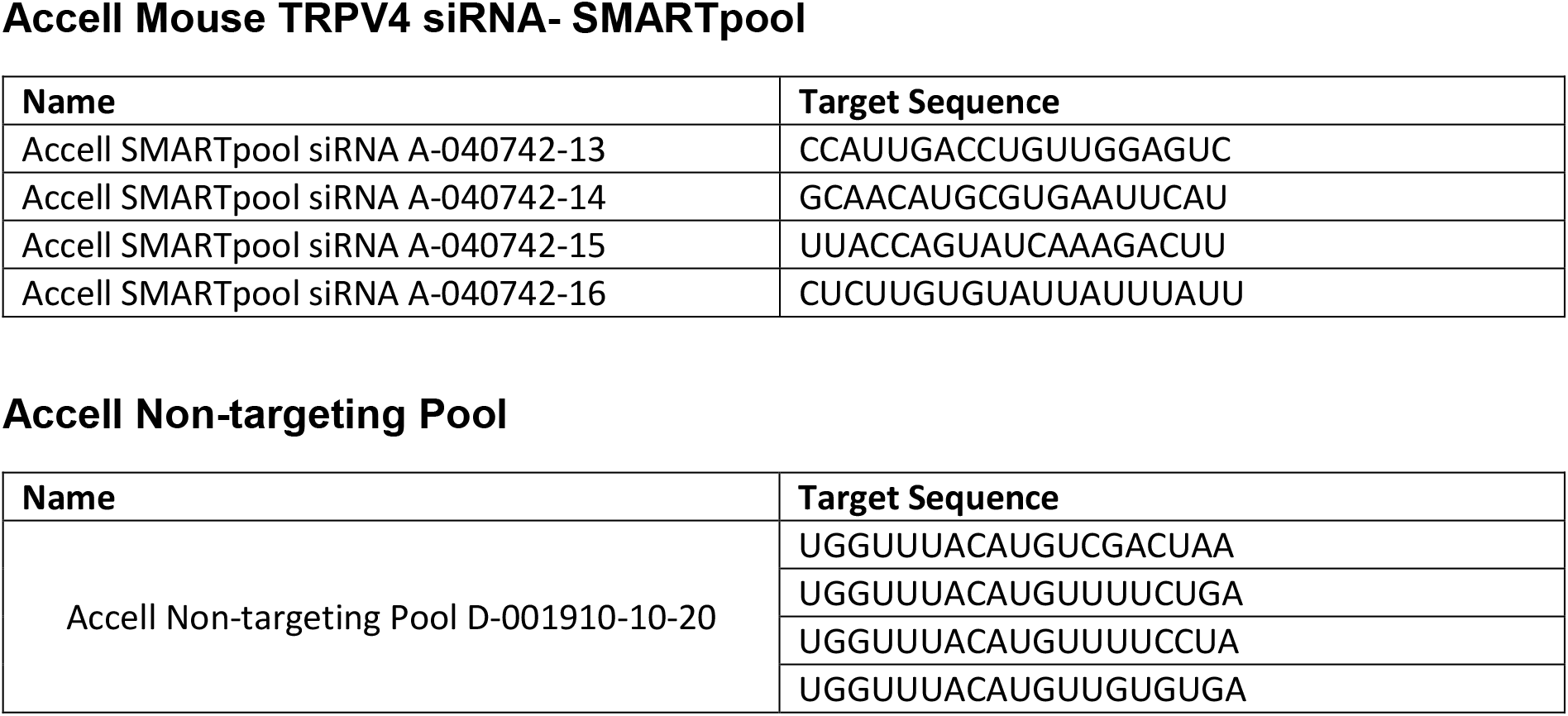
List of oligonucleotides used for nanostring® nCounter expression analysis and RNA sequences for used siRNAs.

### Patch Clamp recordings of ATII cells

Conventional whole-cell recordings were carried out at RT 24 hours after isolation of ATII cells from WT and TRPV4−/− mice. The following bath solution containing 140 mM NaCl, 1.3 mM MgCl2, 2.4 mM CaCl2, 10 mM glucose, 10 mM HEPES (pH 7.4 with NaOH) and resulting in an osmolality of 310 mOsm kg-1 was used for patch-clamp recordings. The pipette solution contained 135 mM CsCl, 2 mM Na-ATP, 1 mM MgCl2, 5 mM EGTA and 10 mM HEPES (pH 7.2 with CsOH), resulting in an osmolality of 296 mOsm kg 1. Patch pipettes made of borosilicate glass (Science Products, Hofheim, Germany) had resistances of 2.2-3.5 MΩ for whole-cell measurements. Data were collected with an EPC10 patch clamp amplifier (HEKA, Lambrecht, Germany) using the Patchmaster software. Current density-voltage relations were obtained before and after application of the TRPV4 activator GSK1016790A (1mM) to the bath solution using voltage ramps from –100 to +100 mV, each lasting 5 s. Data were acquired at a frequency of 40 kHz after filtering at 2.8 kHz. The current density-voltage curves and the current density amplitudes at ±100 mV were extracted at minimal or maximal currents, respectively.

### Western blot analysis

Western Blotting was done as previously described (*Hofmann et al., 2017*). Chemiluminescence was detected in an Odyssey®Fc unit (Licor, Lincoln, NE, USA). Used antibodies and dilutions were: HRP-conjugated anti-β-actin antibody (Sigma A3854HRP, 1:10000); anti-TRPV4 (rabbit, Abcam ab 39260, 1:1000), anti-aquaporine-5 (rabbit, Alomone AQP-005, 1:1000), anti-podoplanin (goat, R&D Systems AF3244, 1:500), secondary anti-goat IgG (whole molecule)-peroxidase (Sigma A5420-1ML, 1:10000) and secondary anti-rabbit IgG peroxidase (POX)-antibody (Sigma A6154, 1:10000).

### Nanostring® nCounter expression analysis

Direct quantification of TRPV4 mRNA in murine lung cells was done as described (*Kannler et al., 2018*). In brief: total RNA from pulmonary murine cells was isolated using the Qia RNeasy Mini Kit (Qiagen, Hilden, Germany). Quantity, purity, and integrity of the RNA samples were controlled by spectrophotometry (NanoQuant, Tecan, Männedorf, Switzerland). Two probes (the reporter and the capture probe) were hybridized to their specific target mRNAs. Then, the target-probe complexes were immobilized in the imaging surface of the nCounter Cartridge by binding of the capture probe. Finally, the sample cartridges were scanned by an automated fluorescence microscope and molecular barcodes (fluorophores contained in the reporter probe) for each specific target were counted. For expression analysis by nCounter NanoString technology, 200 ng total RNA was hybridized with a Nanostring Gene Expression CodeSet and analysed using the nCounter Digital Analyzer. Background correction was performed and normalization was applied using 4 different housekeeping genes (Succinate dehydrogenase subunit A (Sdha), β2-microglobuline, glyceraldehyde 3-phosphate dehydrogenase (GAPDH), β-actin). The DNA sequences used for mRNA expression analysis are summarized in *Table 1*.

### Migration assay

Around 4.4 × 10^6^ ATII cells/well were seeded on a 2 well silicone insert with a 500 µm cell-free gap (ibidi GmbH, Martinsried, Germany) and grown in DMEM (10% FCS, 1% HEPES and 1% penicillin/streptomycin) for 5 days to obtain ATI like cells. Subsequently cells were starved in serum reduced medium (0.1% FCS) for 24 h before insert detachment to create a defined cell-free gap. Images were taken 0, 1, 3, 5, 8, 12 and 24 h after gap creation. Migration was analyzed by measuring the remaining gap width with ImageJ software in 3 pictures per time point and replicate.

### Isolation of nuclear fractions

Isolation of nuclear protein extracts from ATI-like cells after 6 days of culture was performed with a Nuclear Extract Kit according to the manufacturer’s instructions (Active Motif, 40010, La Hulpe, Belgium) as described (Hofmann et al., 2017). In brief, cells were first washed with PBS containing phosphatase inhibitors. Cytoplasmic protein fractions were collected by adding hypotonic lysis buffer and detergent, causing leakage of cytoplasmic proteins into the supernatant. After centrifugation (14.000 × g for 30 s) nuclear protein fractions were obtained by resuspending pellets in detergent-free lysis buffer containing protease inhibitors. NFAT proteins were analyzed by Western blotting as described below using an NFATc1 specific (mouse, SantaCruz Biotechnology, sc-7294, 1:600) antiserum and lamin B1 (rabbit, Thermo Fisher Scientific, PA5-19468, 1:5000) antibodies as loading control. Protein bands were normalized to loading controls and quantified by an Odyssey® Fc unit (Licor, Lincoln, USA).

### Quantification of cell resistance by electrical cell impedance sensing (ECIS)

Resistance changes of ATII to ATI cells was analyzed using an electric cell impedance sensing (ECIS) device (Applied Biophysics, Troy, NY, USA). Freshly isolated epithelial cells were seeded on ECIS culture ware (8W10E+; Applied Biophysics, Troy, NY, USA), which was preincubated with FCS for three hours and connected to the ECIS device. A total of 1 × 10^4^ cells were seeded per chamber and grown at 37°C and 5% CO_2_ in an incubator. Resistance (Ω) was analyzed at 2000 Hz over 160 h.

### Statistics

All our data are based on experiments with mice. According to the 3R (for reduce, refine, replace) rules German and European guidelines aim to reduce animal experiments. Therefore, we stopped all experiments with animals as soon as a clear trend for significance or no significance was observed. Experimental groups were defined by genotype of mice or treatment (ischemia or no ischemia; GSK activator or solvent). All statistical test were performed using GraphPad Prism 7 (GraphPad Software, San Diego, USA).

## Acknowledgements

The authors thank Bettina Braun for excellent technical assistance and the Riken BioResource Center (RBR) as well as the Mutant Mouse Resource and Research Center (MMRRC) for providing the mouse models. This study was funded by the Deutsche Forschungsgemeinschaft (TRR 152, project 16 (AD) and project 22 (GKC), GRK 2338, project 04 (AD) and the German Center for Lung Research (DZL) (AD; MB, NW, TG).

## Funding

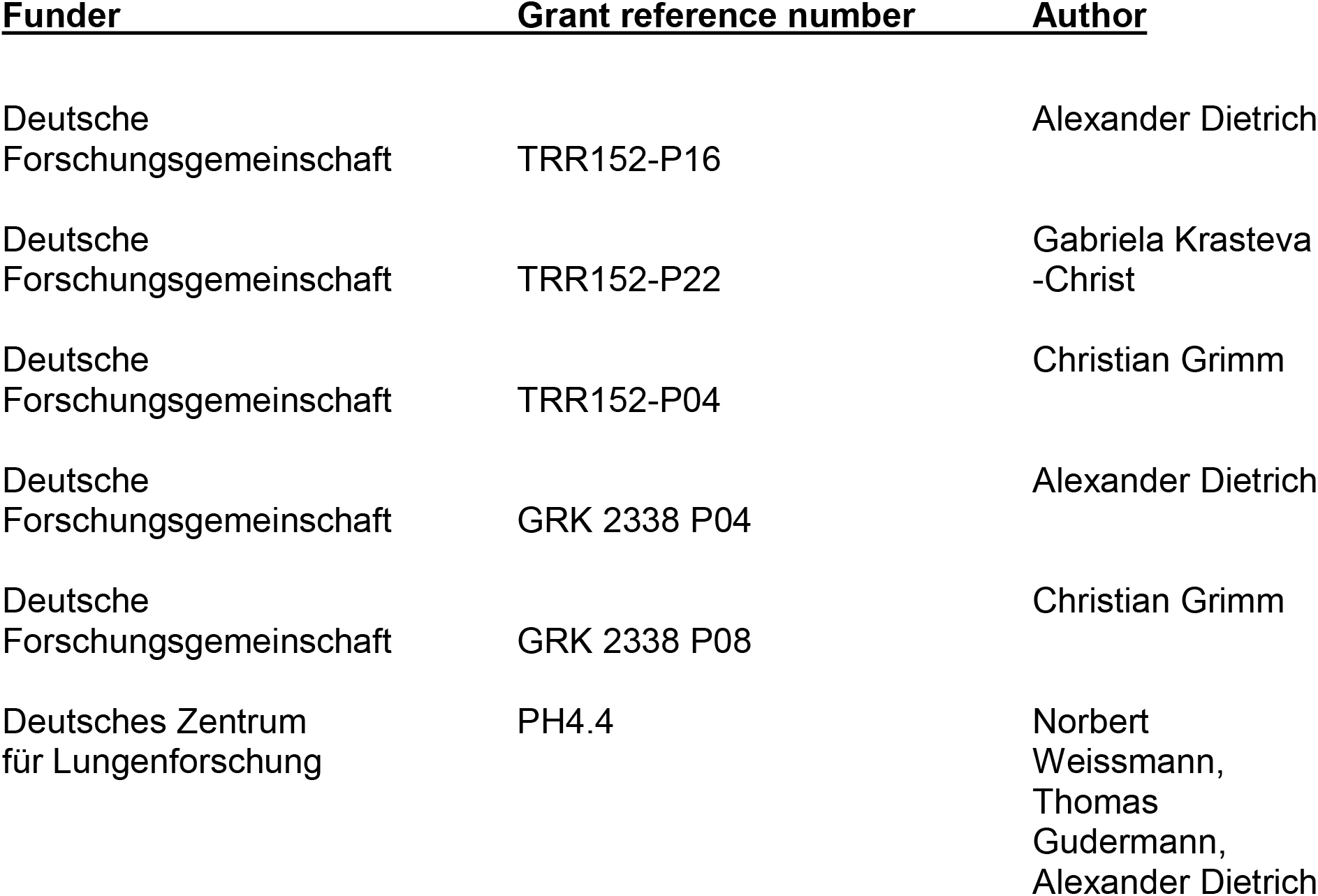

The funders had no role in study design, data collection and interpretation, or the decision to submit the work for publication.

## Author contributions

JW, AD Conception and design, Acquisition of data, Analysis and interpretation of data, Drafting or revising the article; MB, NW, AÖY, JS Contributed essential unpublished data or reagents; Y-KC, MK, GK-C, SR, CG Acquisition of data, Analysis and interpretation of data; TG Drafting or revising the article

## Ethics

Animal experimentation: Experiments involving animals were done in accordance with the EU Animal Welfare Act and were approved by the local councils on animal care (permit No. 55.2.-1-54-2532-7-2015) from Government of Oberbayern, Germany.

---

**Figure 2-figure supplement 1.**
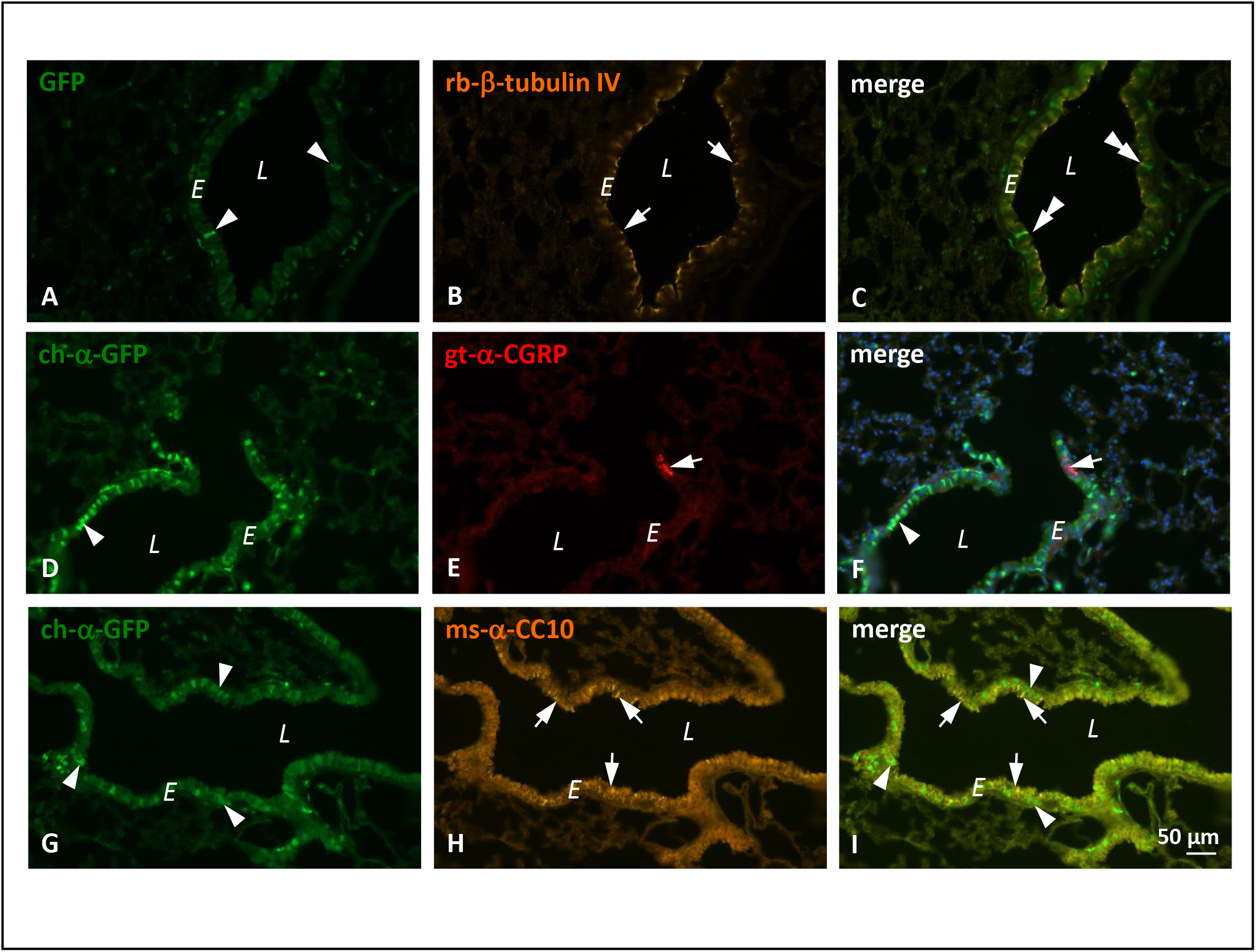
Localization of TRPV4 in mouse lungs using immunohistochemistry. *L* = bronchial lumen, *E* = epithelial layer, scale bars = 50 µm. *A)* Lung cryosections of TRPV4-eGFP reporter mice revealed expression of TRPV4 (arrowheads) in a subpopulation of bronchial epithelial cells. (*B)*. Ciliated cells were labeled for β-tubulin IV (arrows). *C*) A merged view of images shown in *A* and *B*. TRPV4eGFP-positive cells are positive for β tubulin IV (double arrowheads, *A-C*). *D-F*) TRPV4eGFP-fluorescence was enhanced using an anti-GFP-antibody. The same distribution pattern of TRPV4eGFP-immunoreactive cells (arrowheads, D-F) was observed. Neuroepithelial bodies labeled by anti-CGRP-antiserum (arrows, E-F) were not immunoreactive for TRPV4eGFP *(merge, F)*. *(G-I)* TRPV4eGFP-immunoreactive cells (arrowheads, *G*) were not labeled for CC10 (arrows, *H-I*), a marker for club cells.

**Figure 3-figure supplement 1.**
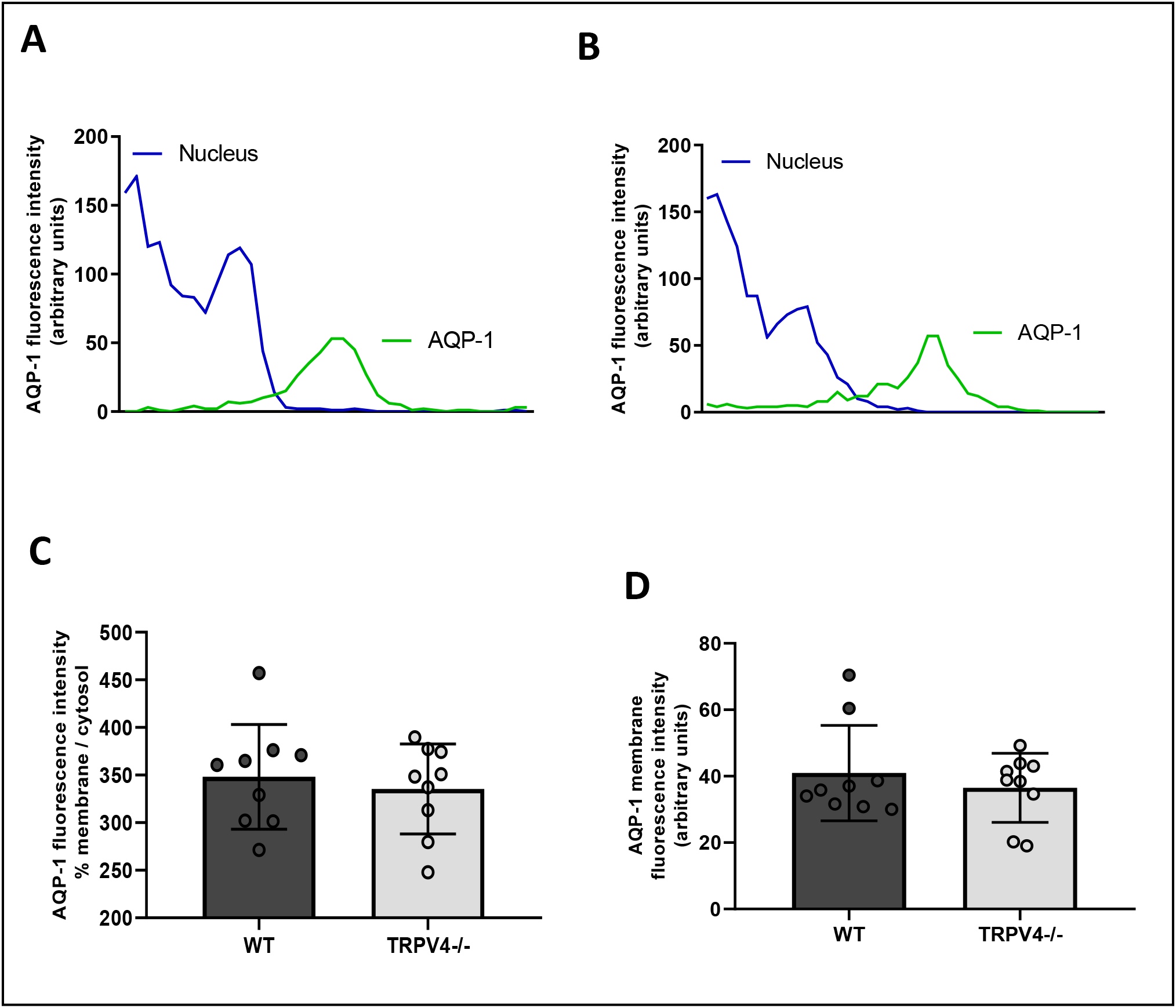
Aquaporin-1 (AQP-1) expression and translocation to the plasma membrane in WT and TRPV4−/− endothelial cells. Representative histograms for the quantification of AQP-1 protein in the plasma membrane of WT (*A*) and TRPV4-deficient endothelial cells (*B*). Summaries of AQP-1 protein expression in plasma membranes (% AQP-1 in membranes (*C*)) and in relation to the cytosol (% AQP-1 membrane/cytosol (*D*)). Data represent means ± SEM from 9 lungs. No significance between means was identified using two tailed unpaired Student’s t-test.

**Figure 5-figure supplement 1.**
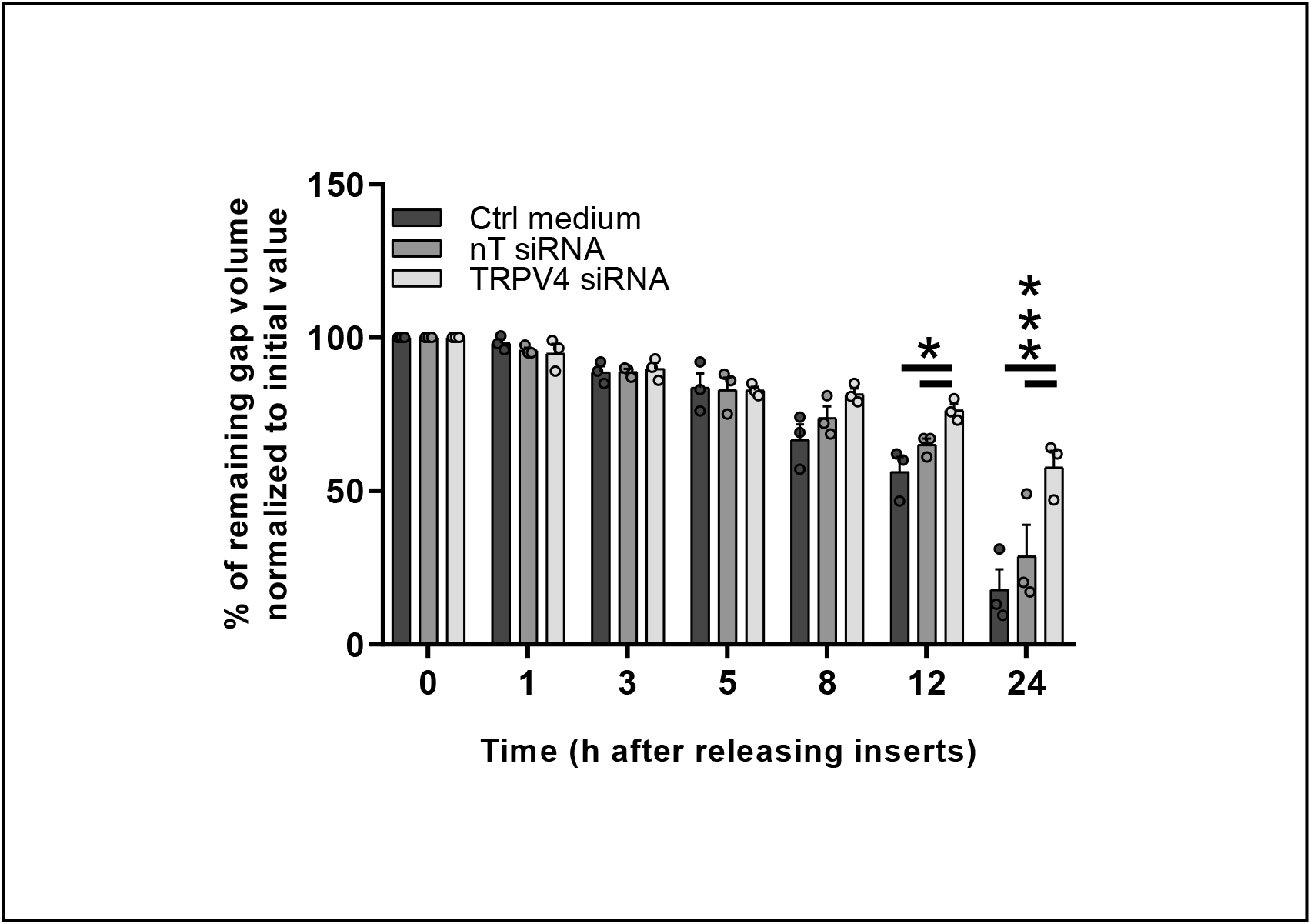
Summary of remaining gap values normalized to initial values quantified in migration assays of WT ATI cells (control (ctrl) medium) as well as ATI cells transfected with siRNA (non-targeting (nT siRNA) or TRPV4-specific siRNAs (TRPV4 siRNA)) after removing inserts at 0, 1, 3, 5, 8, 12 and 24 h. Data represent means ± SEM from 3 independent cell preparations of 5 mice each. Significance between means was analyzed using two way ANOVA and indicated as *** for p<0.001 and * for p<0.05.

